# A prebiotic genetic alphabet as an early Darwinian ancestor for pre-RNA evolution

**DOI:** 10.1101/2023.03.16.532322

**Authors:** Anupam A. Sawant, Sneha Tripathi, Sanjeev Galande, Sudha Rajamani

## Abstract

RNA-based genetic code is thought to be central to life’s emergence due to its dual ability for information transfer and catalysis. Nonetheless, the genetic code of early life was potentially not restricted to canonical genetic alphabets alone. The presence of an extensive repertoire of modified nucleobases in extant biology as ‘signatures of the past’, highlights the relevance of non-canonical alphabets, ably strengthened by experiments demonstrating their ready conversion into nucleosides and nucleotides. All these strongly support a pre-RNA World, wherein informational molecules are posited to have contained alternate genetic alphabets. Nevertheless, understanding pre-RNA molecules’ capacity to acquire emergent function has remained less prevalent. Further, the steps involved in their transition to a canonical RNA World has not been systematically studied in the origins of life framework. In this study, we report the synthesis of a prebiotically relevant genetic alphabet containing the non-canonical nucleobase, barbituric acid. We demonstrate for the first instance the enzymatic incorporation of this prebiotically plausible alphabet (**BaTP**) into an RNA, using proteinaceous T7 RNA polymerase. Pertinently, the incorporation of this genetic alphabet into a baby spinach aptamer did not affect its overall secondary structure, while also allowing it to retain its aptameric function. Furthermore, we demonstrate the faithful transfer of genetic information from pre-RNA-containing barbitudine nucleotides to DNA, using a high-fidelity RNA-dependent DNA polymerase. These findings allude to a putative pathway for the early molecular evolution of the genetic code of extant life.

## Main

Life is thought to have emerged on the early Earth around 3.5b years ago^1–3^. Two approaches that have dictated the origins of life studies include a top-down “biology-centric” approach and a bottom-up “chemistry-inspired” approach^4^. The former is based on phylogenetic analysis to deduce the simplest cellular life form called LUCA (the Last Universal Common Ancestor), which is thought to be a compartmentalized organism with a complex metabolic and catalytic network, along with a sophisticated genetic system^5,6^. The bottom-up approach relies on studying the emergence of life from simpler interacting organic chemicals^7–9^. Understanding the chemical emergence of life through LUCA poses the following two prominent questions, 1) Which biomolecule came about first?^10^ 2) What and how many steps were involved that led to the emergence of the first cellular life? Various models have been proposed, including protein-first^11,12^, metabolism-first^13,14^, and membrane-first models^15,16^, that have been fundamental to understanding life’s emergence. Nonetheless, none have been as attractive as the nucleic acid-first model that centres around the dual ability of RNA to transfer information and perform catalysis^17^. Pertinently, the importance of informational molecules in driving Darwinian evolution further adds weight to this model. The most significant step in RNA-based evolution is the formation of nucleotide monomers under early Earth conditions. This is followed by oligomerization and enrichment of the informational oligomers by adopting secondary structures and acquiring functions, establishing the emergence of an RNA World^18^. Noteworthy is that when the constituents of a nucleoside monomer are allowed to react under prebiotic conditions, it almost always results in the “nucleosidation problem” i.e., the difficulty faced during the condensation of a nucleobase on to the sugar. This problem is further compounded by the low solubility and reactivity of nucleobases (predominantly pyrimidines) in water^19–21^. Therefore, extant nucleobases are not ‘special’ if one aims to study RNA evolution based on just reactivity.

A natural respite to the aforesaid issue is provided by the heterogeneity inherent to the prebiotic soup, as it is a diverse mixture of various heterocycles that have the potential to result in nucleobases^22–24^. In this context, earlier studies have explored the possibility of the formation of nucleo(si)tide monomers (prebiotic genetic alphabets, PGA), using few of these prebiotically relevant heterocycles and sugars under various pertinent early Earth conditions. Strikingly, these prebiotic nucleobases seem to fare better in comparison to extant nucleobases. The above research findings essentially demonstrate that the nucleosidation problem can be circumvented with prebiotic nucleobases^25–30^, along with the prevention of information loss (depurination) that results during RNA oligomerization under putative early Earth geochemical conditions^31^. Most importantly, post glycosylation reactions, PGAs often form analogs that are isomorphic to extant nucleo(si)tides in terms of their potential H-bonding faces^32^. Despite the aforementioned features and the potential of these prebiotic nucleoside/tide surrogates, and their propensity to result in intact informational oligomers (pre-RNAs), a systematic understanding of the ability of PGA containing pre-RNAs, to adopt structures essential to emerging as functional RNA motifs (e.g., aptamers and ribozymes), has remained lacking.

Similarly, in the biology-centric approach, various efforts have been put into creating artificial or semi-synthetic informational polymers to expand the genetic code of extant life^33,34^. The artificial genetic alphabets used in these studies are comprised of modified nucleobases which have been successfully integrated to study their properties as well as characterize the fidelity of information transfer in these systems, using both *in vitro* and *in vivo* approaches^35–37^. Phylogenetic studies of polymerase enzymes show that early (prokaryotic) polymerases promiscuously incorporate non-canonical nucleotides with unnatural or modified nucleobases. These studies raise an important question concerning the genetic code of life and its possible molecular evolution during the genetic transition from ancient life to extant life. Further, the use of several modified nucleobases in extant biology also adds weight to the pre-RNA World conjecture^38^.

In this context, PGAs containing non-canonical nucleobases that are isomorphic to contemporary pyrimidine nucleobases are interesting candidates for studying RNA-based evolution during life’s early stages^39,40^. Studying the formation and emergent functions of early RNA molecules containing non-canonical nucleobases would also be central to understanding the evolution of the genetic system. Against this backdrop, we report the synthesis of one such putative PGA (a triphosphate) that contains barbituric acid nucleobase (Barbitudine triphosphate, **BaTP**). Further, we demonstrate the faithful incorporation of **BaTP** into an RNA using bacterial T7 RNA polymerase. To our knowledge, this is the first report of such incorporation. Despite the incorporation of **BaTPs** into a functional baby Spinach aptamer (bSP) RNA, we show that the folding of the aptameric RNA is not hampered. This emphasizes the suitability of **BaTP**-containing aptamer RNA for acquiring functions which was demonstrated by the binding of this aptamer to its ligand DFHBI.

Furthermore, we also explored the propensity for information transfer from this PGA-containing hybrid RNA to DNA, via reverse transcription and PCR amplification. This is not only important for understanding the faithful transfer of genetic information from barbitudine-modified pre-RNA to DNA, but also has implications for discerning the transition of the genetic code from pre-RNA centric world to the RNA World, followed by the extant World.

### Synthesis of barbitudine, a prebiotic genetic alphabet (PGA) containing barbituric acid

Barbituric acid was recently explored as a prebiotic nucleobase that readily forms nucleoside and nucleotide analogs under prebiotically pertinent conditions^31,40^. Further, nucleotide monophosphate of barbituric acid was shown to undergo oligomerization under strongly acidic conditions at high temperatures. Abiotically formed barbituric acid nucleoside/tide is isomorphic to uridine and pseudo uridine (ubiquitously found in structural RNAs)^41^. Therefore, the nucleoside Barbitudine is an interesting candidate as a prebiotic genetic alphabet to study early RNA evolution. Additionally, Krishnamurthy^42^ and Powner^43^ groups recently explored the route for the prebiotic formation of nucleoside triphosphates, as these are thought to have paved the way from abiotic to biotic transition. Given all these aforementioned studies, we felt encouraged to synthesize Barbitudine triphosphate (**BaTP**), and systematically explore it as a putative pre-RNA World prebiotic genetic alphabet.

We synthesized this barbituric acid-containing PGA by using conventional synthetic organic chemistry. To achieve this, we first allowed the nucleobase BA to react with commercially available acetate donor **1** in the presence of N, O-bis(trimethylsilyl)acetamide (BSA), and TMSOTf as a catalyst (**Fig 1**). The resultant protected ribonucleoside analog **2** was then purified using normal-phase chromatography. Further, this protected ribonucleoside **2** was subjected to deprotection in sodium methoxide and methanol to obtain the ribonucleoside Barbitudine, which was systematically characterized using mass and NMR techniques to confirm its integrity. Since the substrate required for the enzymatic incorporation of this moiety is a triphosphate of barbitudine, the ribonucleoside barbitudine was therefore successfully converted into the barbitudine triphosphate analog (**BaTP**). This was done by a one-pot reaction using POCl_3_, followed by treatment with bis-pyrophosphates in trimethyl phosphate as a solvent (**Fig 1**). The resulting triphosphate **BaTP** was systematically purified, first by anion exchange chromatography, followed by reverse phase chromatography. The final compound was then subjected to HRMS and NMR analysis to confirm its integrity as a triphosphate product (**Fig S1-S4**).

**Figure 1.**
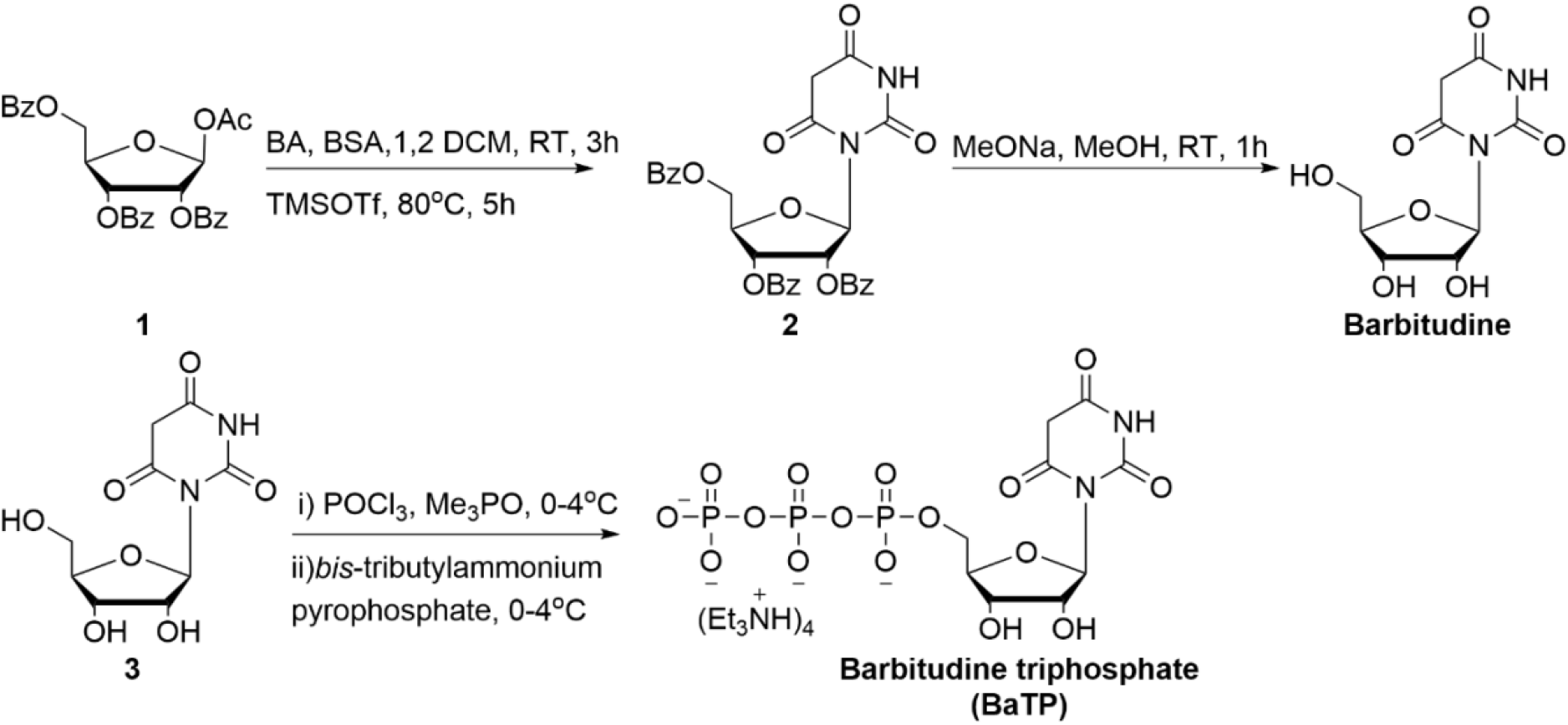
Synthesis of BaTP. Scheme depicting various steps in the chemical synthesis of the prebiotic genetic alphabet Barbitudine and its corresponding triphosphate Barbitudine triphosphate (**BaTP**)^44^.

### Enzymatic incorporation of the BaTP into baby spinach (bSP) RNA aptamer

There are two routes by which the **BaTP** could be incorporated into an RNA to study the emergent properties of pre-RNA motifs; one is using a chemical route (for shorter RNAs) and the other is by using the enzymatic route. Since we aimed to explore the suitability of barbitudine to undergo abiotic to biotic transition, we proceeded with the enzymatic incorporation route as it would provide a ‘natural’ selection pressure for assessing **BaTP’s** incorporation into functional length RNA stretches. Further, this could shed light on the feasibility of molecular evolution of the genetic alphabet, especially during major genetic transitions. Towards this we explored the efficacy of *in vitro* transcription reactions to incorporate **BaTP** into a functional RNA motif^45^. We chose a DNA template (see DNA template design in the Methods section), which typically results in a 43-nt long minimum functional RNA aptamer called the baby spinach (bSP) aptamer, after undergoing successful transcription reaction in the presence of UTP^46^.

This DNA template **D** was now used in a second transcription reaction, wherein , **BaTP** incorporation was assessed by performing reactions in the presence of T7 RNA polymerase, was systematically assessed and the designed DNA template **D** contain ten dA residues in the coding region, which if successful, would guide the incorporation of the monophosphate of **BaTP**, into the RNA transcripts. The transcription reaction was performed in the presence of GTP, CTP, ATP, and UTP/**BaTP**, and the reactions were analysed with 3% agarose gels. Our results showed that the transcription reaction with **BaTP** and template **D** did indeed result in the formation of full-length RNA transcripts that seemed to have incorporated the Barbitudine nucleotide, albeit in moderate yields when compared to the control reaction where the canonical UTP was used (**Fig 2A**, lane 3 vs lane 2). We further assessed the competitive incorporation of **BaTP** versus the canonical UTP by performing an *in vitro* transcription (IVT) reaction in the presence of 1:1 concentration of UTP and **BaTP**. Interestingly, the RNA polymerase incorporated **BaTP** nucleotide(s) effectively in this IVT reaction even in the presence of UTP (**Fig 2B**).

**Figure 1.**
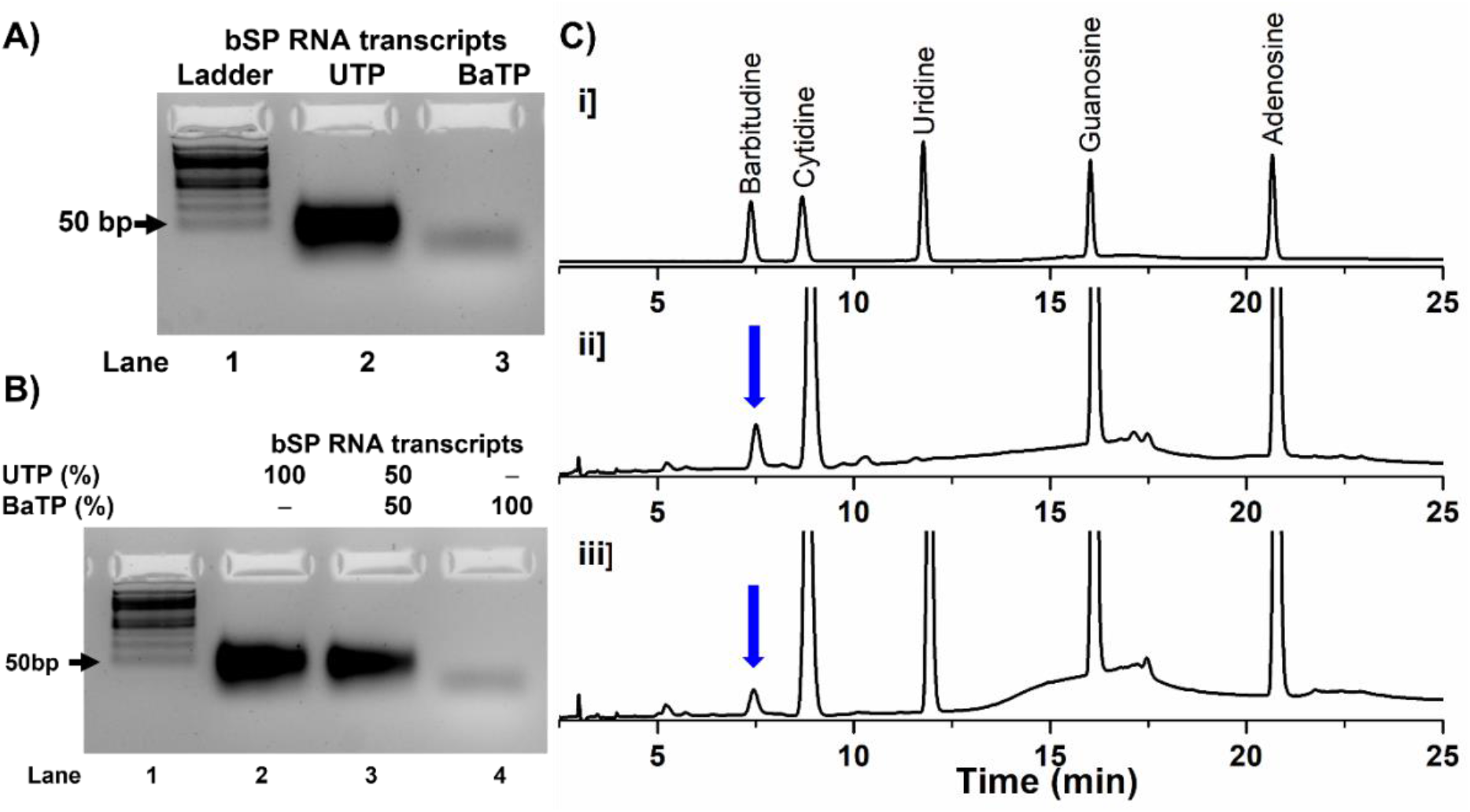
Enzymatic incorporation and digestion of BaTP modified RNA. **A)** 3% agarose gel showing the baby spinach (bSP) RNA product that resulted from *in vitro* transcription (IVT) reactions performed in the presence of UTP and the prebiotic genetic alphabet, **BaTP**. Lane 1, 50 bp ladder. Lane 2 shows the RNA transcripts from the IVT reaction which used the four canonical nucleotides, including UTP. Lane 3 shows the RNA transcripts from the IVT reaction where the UTP was totally replaced with **BaTP**. Comparison of lanes 2 and 3 indicate the formation of full-length RNA transcripts, which should be in the range of 40-50 bp. **B)** The competitive incorporation of natural UTP versus the PGA **BaTP** by T7 RNA polymerase enzyme was assessed by performing a reaction in the presence of an equimolar concentration of UTP and **BaTP** (lane 3). **C)** HPLC chromatograms of ribonucleoside products obtained from the enzymatic digestion of RNA transcripts at 260 nm. (i) Standard mix of natural ribonucleosides and Barbitudine. (ii) RNA digests that were obtained from IVT by transcription reactions carried out in the presence of **BaTP**. (iii) RNA digest obtained from IVT by transcription reaction carried out in the presence of an equimolar concentration of UTP and **BaTP** (band from lane 3 in **Fig 1B**). HPLC details - Mobile phase A: 100 mM TEAA (pH 7.5); mobile phase B: acetonitrile. Flow rate: 1 mL/min. Gradient: 0-10% B in 20 minutes and 10-100% B in 10 minutes.

Next, to confirm the incorporation of barbitudine triphosphate into the bSP RNA aptamer, we performed enzymatic digestion of the RNA obtained after successful transcription reactions with **BaTP** (**Fig 2B**, lane 3), and the reactions performed in the presence of an equimolar concentration of **BaTP** and UTP (**Fig 2B**, Lane 2). The resulting RNA digests were then subjected to HPLC analysis. The HPLC chromatogram for the Barbitudine containing RNA digest did indeed show a peak for barbitudine. This nucleoside eluted at the same retention time as that of the corresponding peak observed in the chromatogram obtained for the mix of nucleoside standards (**Fig 2C**, **i** and **ii** panels respectively). Further, the RNA digest of the competitive incorporation of **BaTP** vs UTP also showed a peak at a similar retention time as barbitudine in the standard mix (**Fig 2C i, ii, iii** panels, respectively). These results further unambiguously ascertained the presence of the genetic alphabet barbitudine, in the full-length bSP RNA transcripts. Additionally, these enzymatic digestion results also highlight the qualitative ability of **BaTP** to compete with canonical UTP, during the transcription reactions.

### Secondary structure analysis of unmodified (UbSP) and modified (BAbSP) RNA aptamers using CD spectroscopy analysis

Circular dichroism (CD) is a very useful technique for characterizing an RNA aptamer that contains G-quadruplex forming sequences. This is pertinent as the G-quadruplex plays a crucial role in the secondary structure formation of bSP RNA aptamer, which turns impinges on its (functional) capacity to bind its ligand. Therefore, for CD analysis, both 5 μM unmodified (UbSP) and 5 μM modified (BAbSP) RNAs were prepared in the presence of a sensor buffer that contained 10 mM Tris-HCl, 100 mM KCl, 10 mM MgCl2 at pH 7.4 (see Methods for sample preparation). The K^+^ ions in the sensor buffer trigger the G-quadruplex formation and the typical signature of a G-quadruplex was readily observed in the unmodified RNA aptamer (UbSP) as shown in **Fig 3C** (red trace). The CD spectrum of the UbSP aptamer also revealed a positive band at 270 nm, and smaller negative peaks at 238 nm and 215 nm, indicating the formation of a mixture of the hybrid, parallel/anti-parallel G-quadruplex, and is consistent with the literature^47,48^. Interestingly, a similar CD result was obtained for the modified bSP RNA aptamer as well (BAbSP, **Fig 3C**, blue trace), in which 10 uridine residues were replaced by a prebiotically pertinent genetic alphabet, barbitudine. Although the ellipticity values on Y-axis indicate only a marginal destabilizing effect due to the barbitudine residues on the modified BAbSP aptamer, the overall secondary structure was still in good agreement with the control unmodified UbSP aptamer.

**Figure 3.**
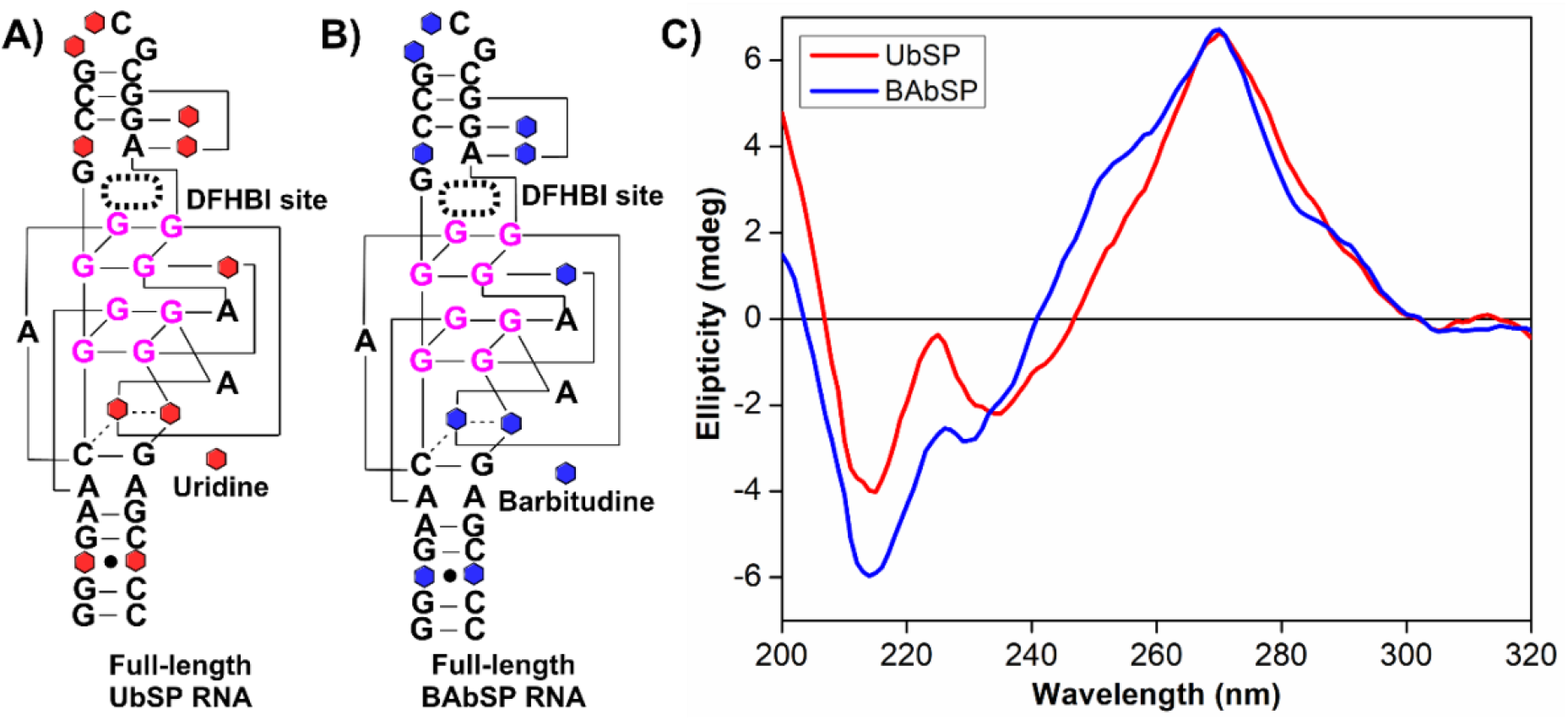
Secondary structure of UbSP and BaSP aptamer RNAs. CD spectra of 5 μM unmodified UbSP (**A** and **C**, coloured red) and modified BAbSP (**B** and **C**, coloured blue) RNA, respectively. Both RNA aptamer oligos were prepared in a sensor buffer containing 10 mM Tris-HCl, 100 mM KCl, 10 mM MgCl_2_, and at pH 7.4.

### Biophysical analysis of modified baby spinach RNA aptamer (bSP) using fluorescence spectroscopy

To act as a Darwinian ancestor, the prebiotic genetic alphabet containing aptamer must demonstrate the ability to fold into the requisite secondary structure that results in emergent functions, like ligand binding in the case of an aptamer, or facilitate catalysis. To test this hypothesis, we first designed and synthesized the basic functional aptamer RNA that contained the prebiotic non-canonical genetic alphabet, **BaTP**. CD analysis confirmed the ability of this modified RNA aptamer BAbSP to adopt a functionally active secondary structure. To further affirm whether the BAbSP aptamer was indeed in an active form, we performed fluorescence analysis of this aptamer in the presence of the ligand, DFHBI (3,5-Difluoro-4-hydroxyphenyl) methylene]-3,5-dihydro-2,3-dimethyl-4H-Imidazole-4-one). If the modified BAbSP RNA shows fluorescence enhancement after binding to the DFHBI ligand, it will reconfirm, firstly, the ability of the barbitudine modified RNA to adopt the correct secondary structure, and, secondly and most importantly, the ability to acquire function, which in this case is that of an aptameric function of ligand binding.

Therefore, we tested the binding of both the control and modified versions of the bSP RNA aptamers, to the non-fluorescent dye DFHBI. This ligand only shows fluorescent signal enhancement when it is bound to the target aptamer RNA. We initially prepared both unmodified UbSP (5 μM) and the modified BAbSP (5 μM) aptamer RNAs in a sensor buffer, using a similar protocol used for the sample preparation during CD analysis. To these preformed aptamer RNAs, 10 μM of the DFHBI ligand was added and the samples were incubated in the dark for 2h. All the respective control samples were also prepared similarly. After the incubation period, the samples were analysed using fluorescence spectroscopy. RNA samples (UbSP and BAbSP) in the absence of the ligand DFHBI (L), did not show any detectable fluorescence (**Fig 4C**, red and blue traces, respectively). Further, the DFHBI alone control showed very weak fluorescence in the absence of the target aptamer RNA (**Fig 4C**, black trace). However, both the aptamer RNAs (UbSP and BAbSP) showed a significant increase in fluorescence intensity when incubated with the DFHBI ligand. The unmodified aptamer RNA, UbSP displayed a 5-fold increase (**Fig 4C**, pink trace), while the modified aptamer RNA, BAbSP showed a nearly 4-fold increase (**Fig 4C**, green trace), in the fluorescence intensity. Overall, the fluorescence analysis indicated that the incorporation of this prebiotic genetic alphabet did result in a variant that folded correctly and demonstrated the ability for functional aptamer binding.

**Figure 4.**
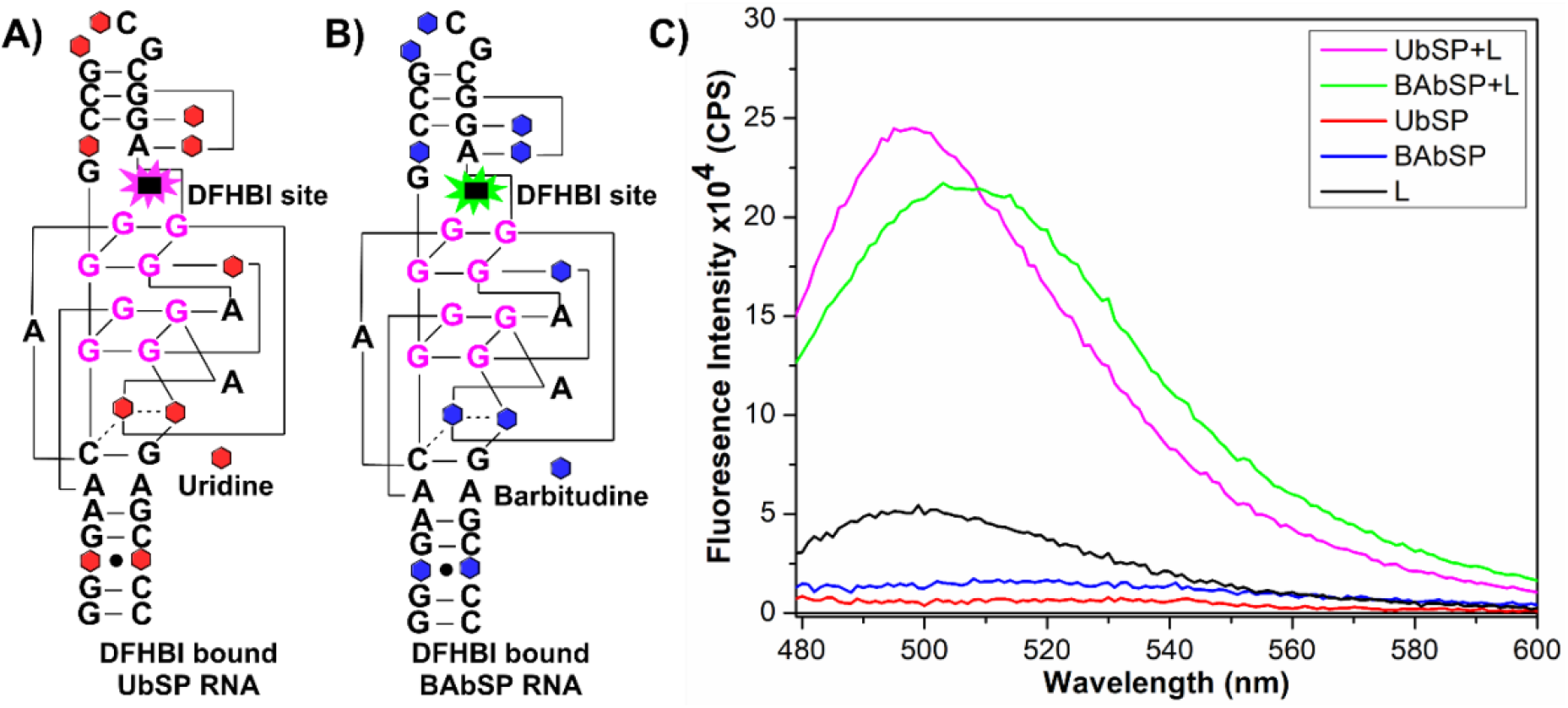
Fluorescence analysis of UbSP and BAbSP aptamer RNAs. Both UbSP and BAbSP aptamer RNAs (5 μM) were prepared in the sensor buffer containing 10 mM Tris-HCl, 100 mM KCl, 10 mM MgCl_2_, and at pH 7.4. To these RNAs, 10 μM of the DFHBI ligand (L) was added and the mixture was further incubated for 2h in the dark. An evident increase in fluorescence intensity was observed for UbSP (pink trace) and BAbSP (green trace) RNAs in the presence of DFHBI. The control samples of only UbSP (red trace), only BAbSP (blue trace), and DFHBI (black trace) alone, did not show any fluorescence. (Excitation at 460 nm and emission at 500-505 nm)

### Information transfer from RNA containing barbitudine to DNA through reverse transcription and PCR amplification

Given the success with T7 polymerase incorporation of **BaTP**, we therefore wished to evaluate the propensity of other extant enzymes for their ablity to use **BaTP** effectively. PCR amplification and sequencing are the essential next steps in this process that would allow to measure the fidelity of canonical enzymes for tolerating the incorporation of a PGA-like **BaTP**. Before doing this, we aimed to first confirm the incorporation of **BaTP** into a longer RNA. Towards this, a ~700 nt long mCherry mRNA was transcribed by transcription reaction using a DNA template, in the presence of UTP/**BaTP** nucleotides and T7 RNA polymerase. To confirm the **BaTP** incorporation, we performed the enzymatic digestion of mCherry mRNA and subjected this RNA digest to HPLC analysis. The HPLC profile clearly showed the incorporation of barbitudine into the modified mCherry mRNA (**Fig S5**). After this confirmation, high-density RNA transcripts containing barbitudine were subjected to RNase-free DNase treatment for 1h at 37°C. The RNA transcripts were purified from DNase-treated reactions and 1ug of RNA sample was subjected to reverse transcription by using Superscript-III™ reverse transcriptase and random hexamer reverse primers to generate the complementary DNA (cDNA). Further, the cDNAs generated from the respective RNAs were subjected to PCR amplification using high-fidelity *Pfu* DNA polymerase in the presence of appropriate forward and reverse primers. We used the following controls i.e the minus RT (reaction without the RT enzyme) control and the NTC (non-template control) sample to confirm the successful reverse transcription of the RNA transcripts.

PCR products obtained after amplification reactions were directly resolved on 1.5 % agarose gel to confirm cDNA synthesis (**Fig. 5A**). Importantly, all the positive reverse transcription reactions (+RT samples), showed the amplicon band. Importantly, the absence of a band from the negative control reaction (-RT) also further confirmed the successful conversion of barbitudine modified RNA transcripts into cDNA, in these reverse transcription reactions. Further, to evaluate the fidelity of the transcription and reverse transcription reactions, PCR amplicons obtained from the amplification of the respective cDNA products were sequenced. The sequence analysis of the PCR products showed 99% of sequence indentities with the template DNA sequence used for the incorporation of **BaTP** into RNA by *in vitro* transcription reaction and most imporatntly, sequence blast clearly showed that no misincorporation was observed while incorporating thymidine against adenosine residue in the cDNA template **(Fig. S7 and S9).** Taken together, these results confirmed that **BaTP** is efficiently incorporated into RNA transcripts (against cognate deoxyadenosine), which are effectively recognized and copied by reverse transcriptase to produce the corresponding cDNA (against barbitudine pre-RNA template). This ability of **BaTP** to preserve the sequence information in a series of steps namely, transcription, reverse transcription, and amplification, would have been beneficial for faithful information transfer in an early RNA World, and its subsequent transition to the RNA-DNA-protein world.

**Figure 5.**
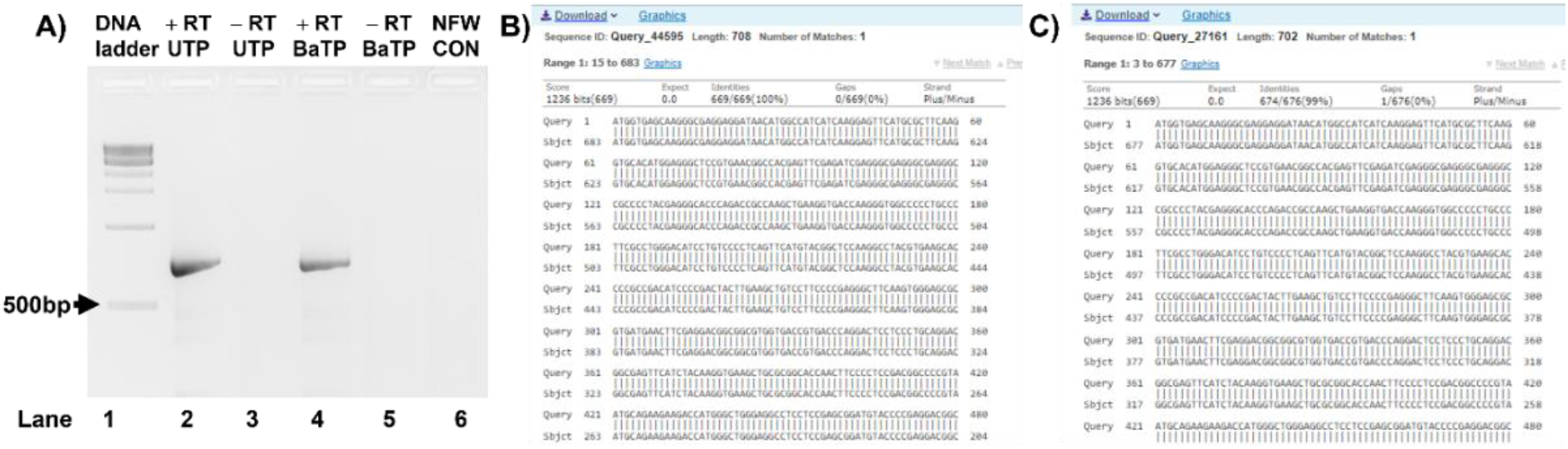
RTPCR and sequence alignment results for unmodied and modified mCherry RNA. **A**) Gel picture depicting PCR amplicons obtained from the various cDNA. PCR reactions were performed with cDNA that were generated after successful reverse transcription of control RNA containing UTP, and modified RNA containing **BaTP** (+RT), respectively. DNase-treated RNA was used as negative reaction control (-RT). Representative sequence alignment (BLAST) of PCR amplified DNA products obtained from cDNA, which was in turn obtained by reverse transcribing **B**) unmodified RNA and **C**) modified RNA containing barbitudine. The sequencing data of the PCR product showed 99% sequence indentities with the template DNA sequence used for the incorporation of **BaTP** into RNA, using *in vitro* transcription reactions.

## Conclusion

One of the fundamental hallmarks of life is its tremendous capacity for evolution which manifests at various levels. Successful attempts to expand the genetic code of extant life to create artificial or semi-synthetic life, confirm that evolution can be tinkered with by externally facilitating changes happening at the molecular levels. The effective proliferation of life requires molecules that can store and propagate information, and perform catalysis. It is widely accepted that informational molecules of life (RNA and also DNA, were) were first formed abiotically, following which molecules with the ability to acquire functions (like aptamers and ribozymes) emerged and were selected for thereafter. Nonetheless, the abiotic emergence of a molecule like RNA, readily from a heterogenous prebiotic soup, is a non-trivial process even under the best of circumstances. Given this, the systematic understanding of how primitive informational polymers of a pre-RNA World emerged and functioned is fundamental to understanding how biotic life evolved from abiotic life. Against this backdrop, we report the synthesis and also enzymatic incorporation of the prebiotically pertinent genetic alphabet **BaTP**, using prokaryotic proteinaceous T7 RNA polymerase. To our knowledge, such enzymatic incorporation of **BaTP** has been reported for the first time.

We observed that **BaTP** can be effectively incorporated by this old-world RNA polymerase to result in a full-length RNA product. Additionally, **BaTP** also competes with its canonical counterpart UTP during the transcription reaction. However, qualitatively speaking, the T7 polymerase preference, understandably, seems to be more towards the canonical alphabet than the non-canonical one. This could be an indication of the early selection pressure on such prebiotic nucleotides, however we need more systematic quantitative studies to affirm this. Further, we also demonstrated the ability of these putative pre-RNA aptamers to adopt secondary structures using CD analysis. This result indicates a similar H-bonding pattern between uridine and barbitudine, as the CD profiles of the unmodified and modified RNA aptamers does not indicate any severe perturbation to its secondary structure. Finally, these results were further confirmed using fluorescence analysis wherein we showed that the aptamer obtained by incorporation of a prebiotically pertinent nucleotide, does not hamper even the function of the aptamer RNA. It showed effective binding to its ligand (DFHBI), which was comparable in fluorescence (enhancement) as the control unmodified aptamer RNA.

Collectively, these results indicate that the prebiotic genetic alphabet barbitudine has the potential to act as an early Darwinian ancestor of modern pyrimidine nucleotides. Further, we have systematically undertaken experiments showing the ability of information transfer of barbitudine and the fidelity associated with it.We found that the RNA that contained barbitudine could act as a template for the synthesis of complementary DNA in the presence of high-fidelity reverse transcriptase. Interestingly, the cDNA could also be further amplified using PCR reaction to yield full-length DNA product, similar to the one used in the original transcription reactions. These are very important observations, which highlight the potential of pre-RNA-containing barbitudine to be suitable for processes like transcription (formation of RNA on the DNA template), reverse transcription (pre-RNA acts as a template for DNA synthesis) that happens central dogma of extant biology. We are currently beginning to explore the ability of mRNAs that have these PGAs to undergo protein synthesis, and are also studying nucleotide metabolism using prebiotic genetic alphabets, both *in vitro* and *in cellulo.*

## Methods

### Materials

Trimethylsilyl trifluoromethane sulfonate, O-bis(trimethylsilyl)acetamide, Barbituric acid, 1-O-Acetyl-2,3,5-tri-O-benzoyl-beta-D-ribofuranose, Sodium methoxide and DOWEX 50 x 8 H^+^ resin was all procured from Sigma Aldrich. DNA oligonucleotides were purchased from Eurofins, Inc. and were used without further purification. T7 RNA polymerase, ribonuclease inhibitor (RiboLock), NTPs, RNase A and RNase T1 were obtained from Fermentas Life Science. Calf intestinal alkaline phosphatase (CIP) and snake venom phosphodiesterase I were procured from Invitrogen and Sigma-Aldrich, respectively. Chemicals for preparing buffer solutions were purchased from Sigma-Aldrich (BioUltra grade). Autoclaved water was used in all the biochemical reactions and for HPLC analyses HPLC grade solvents were used.

### Instrumentation

NMR spectra of small molecules were recorded on 400 MHz Jeol ECS-400 and Bruker AVANCE III HD ASCEND 400 MHz spectrometers and processed using Mnova software from Mestrelab Research. Mass analysis was performed using the ESI-MS Waters Synapt G2-Si Mass Spectrometry instrument. HPLC analysis was done using Prominence UFLC Next Generation High Throughput-Shimadzu HPLC using reverse phase shim-pack GIST C18 column (250 x 4.6 mm, 5 microns). The absorption spectra were recorded on a UV-2600 Shimadzu spectrophotometer. Steady-state and time-resolved fluorescence spectra of the control and modified spinach aptamers were recorded on a Fluoromax-4 spectrophotometer (Horiba Scientific). CD analysis was performed on a JASCO J-815 CD spectrometer.

### Synthesis of 6-oxo uridine-5’-triphosphate (BaTP)

An ice-cold solution of the prebiotic ribonucleoside Barbitudine (50 mg, 0.27 mmol, 1.0 equiv.) in trimethyl phosphate (1 mL) was slowly added to freshly distilled POCl_3_ (37 μL, 0.40 mmol, 1.5 equiv.). The reaction mixture was stirred for 15 h at ~4 °C. A solution of bis-tetrabutylammonium pyrophosphate (0.5 M in DMF, 2.75 mL, 5.1 equiv.) and tributylamine (0.65 mL, 2.75 mmol, 10 equiv.) was rapidly added under ice-cold conditions. The reaction was quenched after 30 min with 1 M triethylammonium bicarbonate buffer (TEAB, pH 7.6, 15 mL) and was extracted with ethyl acetate (2 × 10 mL). The aqueous layer was evaporated and the residue was purified, first on a DEAE Sephadex-A25 anion exchange column (10 mM–1 M TEAB buffer, pH 7.6), followed by reversed-phase flash column chromatography (C18 RediSep Rf, 0–40% acetonitrile in 100 mM triethylammonium acetate buffer, pH 7.5, 50 min). Appropriate fractions were lyophilized to separate out the desired triphosphate product **BaTP** as a tetra triethylammonium salt (42 mg, 16%). ^1^H NMR (400 MHz, D_2_O) δ 6.15 (d, J = 2.8 Hz, 1H), 4.68 (dd, J = 6.3, 3.1 Hz, 2H), 4.42 (t, J = 6.7 Hz, 1H), 4.22 (dd, J = 8.5, 5.7 Hz, 1H), 4.08 – 4.01 (m, 2H). ^13^C NMR (101 MHz, D2O) δ 179.73 (s), 166.67 (s), 87.38 (s), 81.15 (s), 71.72 (s), 69.46 (s), 65.84 (s), 58.66 (s). ^31^P NMR (162 MHz, D_2_O) δ - 10.58 (dd, J = 49.6, 19.8 Hz), −22.83 (s). HRMS: (m/z) Calculated/expected mass for C_9_H_14_N_2_O_16_P_3_ [M-H]- = 498.9556, Observed mass = 498.9553.

### *In vitro* transcription with DNA template in the presence of BaTP

The transcription reactions were carried out in 40 mM Tris-HCl buffer (pH 7.9) containing 200 ng of the DNA template (see DNA template design below), 1 mM of GTP, CTP, ATP, UTP/**BaTP**, 10 mM MgCl2, 10 mM NaCl, 10 mM of dithiothreitol (DTT), 2 mM spermidine, 1 U/μL RNase inhibitor (Riboblock) and 40 units of T7 RNA polymerase, in a total volume of 25 μL. Samples were incubated overnight at 37 °C. the RNA products were purified by phenol-chloroform and the RNA pellet formed from each reaction was washed with 70% aqueous ethanol, dried, and dissolved in 20 μL of nuclease-free water. Control and modified transcripts obtained after transcription reactions were resolved and detected by 2 % agarose gel containing EtBr. Further, the presence of the Barbitudine (**3**) in the RNA transcript was confirmed by enzymatic digestion followed by HPLC analysis of ribonucleoside products obtained from the digestion reaction (For experimental details see enzymatic digestion in method section)

Details of DNA template design: Region underlined and highlighted in cyan depicts the binding site for the reverse primer. Region underlined and highlighted in yellow depicts the binding site for the forward primer, while the coding region is highlighted in cyan color.

<u>5’TAA TAC GAC TCA CTA TAG</u> GGT GAA GGA CGG GTC CGT TCG CGT TGA <u>GTA GAG TGT GAG CTC C 3’</u>

### Enzymatic digestion of BAbSP

~Four nmol of the modified oligoribonucleotide transcripts that were obtained after transcription reaction carried out either in the presence of **BaTP**, or with equimolar concentration of **BaTP**: UTP were treated separately with snake venom phosphodiesterase I (0.01 U), calf intestinal alkaline phosphatase (10 μL, 1 U/μL) and RNase A (0.25 μg), in a total volume of 100 μL in 50 mM Tris-HCl buffer (pH 8.5, 40 mM MgCl2, 0.1 mM EDTA) for 12 h at 37 °C. After this period, RNase T1 (0.2 U/μL) was added, and the samples were incubated for another 4 h at 37 °C. The ribonucleoside mixture obtained from the digest was analysed by reversed-phase HPLC using the Shimadzu Shim-pack GIST C18 column (250 x 4.6 mm, 5 microns) at 260 nm. Mobile phase A: 50 mM triethylammonium acetate buffer (pH 7.5), mobile phase B: acetonitrile. Flow rate: 1 mL/min. Gradient: 0-10% B in 20 minutes and 10–100% B in 10 minutes.

### CD analysis of control (UbSP) and modified (BAbSP) aptamer RNA

For CD analysis, solutions of 5 μM control Baby Spinach aptamer RNA (UbSP) and modified aptamer RNA (BAbSP) were prepared in sensor buffer containing 10 mM Tris pH 7.5, 50 mM KCl and 5 mM MgCl_2_, in nuclease-free water. The samples were vortexed and spun down, and annealed for 4 min at 90 °C in a heating block. Samples were further allowed to come to RT slowly on the heating block. CD analyses was carried out right after that and the CD spectra were recorded on a Jasco J-815 spectrometer using a standard quartz cuvette with a 1-cm path length. Scans from 200 to 320 nm were acquired with a scanning speed of 100 nm/min, a step of 1 nm and a bandwidth of 1 nm. A positive band at 264 nm confirmed the formation of a G-quadplex binding pocket, as previously reported by Dasgupta *et. al.* for G-quadruplex-containing RNAs, including Spinach derivatives^48^. Identical CD spectra obtained for the modified Baby Spinach RNA BAbSP confirmed the formation and the intactness of the G-quadruplex binding pocket.

### Fluorescence analysis of control (UbSP) and modified (BAbSP) aptamer RNA

RNA samples (5 μM) were prepared for fluorescence analysis using a similar protocol as described for CD analysis. In addition, a DFHBI ligand (10 μM) was added to the RNA samples, UbSP and BAbSP respectively. The control RNA samples and the DFHBI control were also prepared in an exactly similar fashion. Before fluorescence analysis, RNA samples were incubated in the dark for 2h. Following the incubation, fluorescence analysis was recorded on a Fluoromax-4 spectrophotometer (Horiba Scientific) using excitation at 460 nm and emission at 500-505 nm. The slit width used for excitation and emission was 8/8 respectively. All the respective controls were also analysed using similar instrumental parameters.

### Reverse transcription and PCR amplification of barbitudine-modified mCherry mRNA

One mg of mCherry mRNA was taken for cDNA synthesis using Superscript III first strand synthesis kit (Invitrogen catalog # 18080051). The cDNA synthesis was done as per the manufacturer’s recommended protocol. Briefly, the RNA, random hexamer and dNTPs were incubated at 65 °C for 5 min, and immediately put on ice (4°C) for another 5 min. The 5X first strand synthesis buffer was added to the above mix, along with 0.1 M DTT, RNAse Inhibitor, and the Superscript III enzyme. The above mixture was incubated at 25 °C for 5 min, 50 °C for 30 min, and 70 °C for 15 min. cDNA validation was performed by using HiFi PFU enzyme (dxbidt catalog # R1220) as per the PCR protocol. Briefly, the PFU master mix (2X) was added to the forward/reverse primer (see details below) and the cDNA template, and the PCR was set up as per the protocol given.

### Details of mCherry DNA template design: Region underlined depicts the binding site for the reverse primer and the forward primer

<u>ATGGTGAGCAAGGGCGAGGAG</u>GATAACATGGCCATCATCAAGGAGTTCATGCGCT TCAAGGTGCACATGGAGGGCTCCGTGAACGGCCACGAGTTCGAGATCGAGGGCGAGGGCGAGGGCCGCCCCTACGAGGGCACCCAGACCGCCAAGCTGAAGGTGACCAAGGGTGGCCCCCTGCCCTTCGCCTGGGACATCCTGTCCCCTCAGTTCATGTACGGCTCCAAGGCCTACGTGAAGCACCCCGCCGACATCCCCGACTACTTGAAGCTGTCCTTCCCCGAGGGCTTCAAGTGGGAGCGCGTGATGAACTTCGAGGACGGCGGCGTGGTGACCGTGACCCAGGACTCCTCCCTGCAGGACGGCGAGTTCATCTACAAGGTGAAGCTGCGCGGCACCAACTTCCCCTCCGACGGCCCCGTAATGCAGAAGAAGACCATGGGCTGGGAGGCCTCCTCCGAGCGGATGTACCCCGAGGACGGCGCCCTGAAGGGCGAGATCAAGCAGAGGCTGAAGCTGAAGGACGGCGGCCACTACGACGCTGAGGTCAAGACCACCTACAAGGCCAAGAAGCCCGTGCAGCTGCCCGGCGCCTACAACGTCAACATCAAGTTGGACATCACCTCCCACAACGAGGACTACACCATCGTGGAACAGT ACGAACGCGCCGAGGGCCGCCACTCCACCGGC<u>GGCATGGACGAGCTGTACAAGT AG</u>

### Forward primer

5 ‘ TAATACGACTCACTATAGGGGATGGTGAGCAAGGGCGAGGAG3’

### Reverse primer

5’ CTTGTACAGCTCGTCCATGCC3’

## Acknowledgements

AAS thanks DBT-RA, DBT, Government of India for funding support. SR thanks SERB, DST [CRG/2021/001851] and DBT, Government of India [BT/PR19201/BRB/10/1532/2016] for funding. SG is a recipient of the JC Bose Fellowship (JCB/2019/000013) from the Science and Engineering Research Board, Government of India. AAS and SR wish to extend thanks to collaborators SG and ST for their support with molecular biology-related resources. SR and AAS express their special gratitude to Dr. Srivatsan’s & Dr. Hotha’s lab for help with certain reagents and providing workspace access for synthesis work.

## Author contributions

AAS and SR conceived the project and designed the experiments. AAS performed the chemical synthesis of the nucleoside triphosphate (**BaTP**) and performed all the biophysical experiments and analysis, and the enzymatic digestion assays and related experiments. ST performed *in vitro* transcription reactions, RT-PCR reactions and gel analysis with crucial inputs from SG. AAS and SR wrote the manuscript with critical inputs from ST and SG.

## Notes

### Competing Interest Statement

The authors have declared no competing interest.

