## Supplementary material for "A prebiotic genetic alphabet as an early Darwinian ancestor for pre-RNA evolution": SI info


| Content | Page |
|---|---|
| 1. **NMR and HRMS spectra** | |
| **Fig S1** 1H-NMR of prebiotic genetic alphabet BaTP in $D_2O$ | 2 |
| **Fig S2** 13C-NMR of prebiotic genetic alphabet BaTP in $D_2O$ | 2 |
| **Fig S3** 31P-NMR of prebiotic genetic alphabet BaTP in $D_2O$ | 3 |
| **Fig S4** HRMS of prebiotic genetic alphabet BaTP in $D_2O$ | 3 |
| 2. **Enzymatic digestion of barbitudine-modified mCherry RNA** | |
| **Fig S5** HPLC chromatograms of ribonucleoside products obtained from enzymatic digestion of RNA transcripts at 260 nm. | 4 |
| **3. PCR product Sequencing chromatogram and sequence alignment with DNA template** | |
| **Fig S6** Representative sequencing chromatogram of PCR amplified DNA products obtained from cDNA, which was in turn obtained by reverse transcribing control unmodified RNA. | 5 |
| **Fig S7** Representative sequencing chromatogram of PCR amplified DNA products obtained from cDNA, which was in turn obtained by reverse transcribing barbitudine modified RNA. | 6 |
| **Fig S8** Representative sequence alignment (BLAST) of PCR amplified DNA product obtained from cDNA, which was in turn obtained by reverse transcribing unmodified RNA. | 7 |
| **Fig S9** Representative sequence alignment (BLAST) of PCR amplified DNA product obtained from cDNA, which was in turn obtained by reverse transcribing barbitudine modified RNA. | 8 |

### 1. NMR and HRMS Spectra

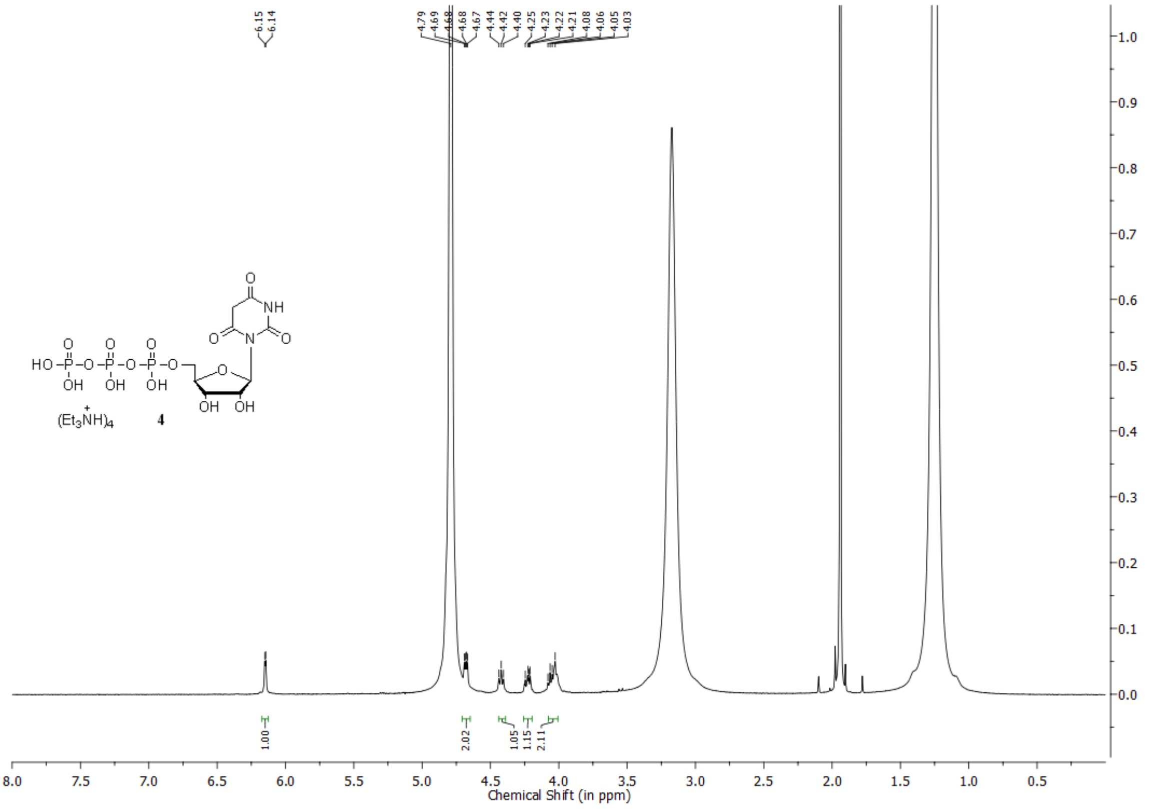

**Fig S1** 1H-NMR of prebiotic genetic alphabet BaTP in $D_2O$

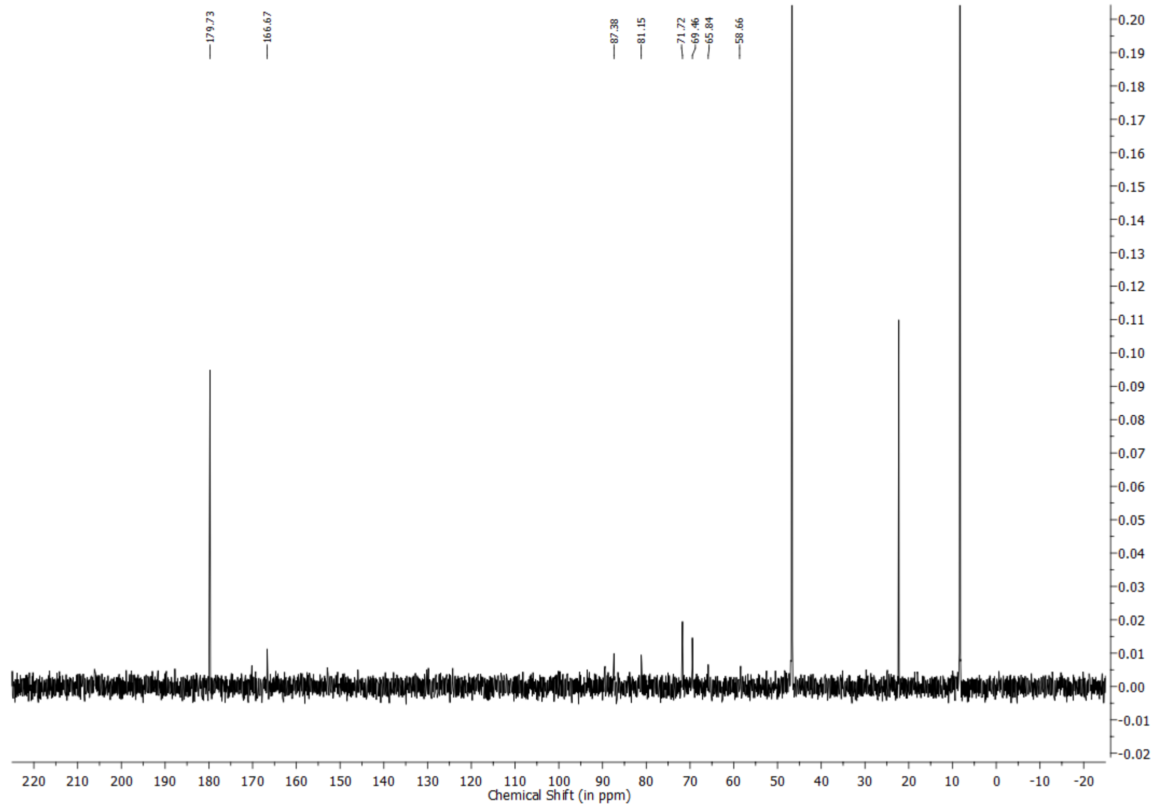

**Fig S2** 13C-NMR of prebiotic genetic alphabet BaTP in $D_2O$

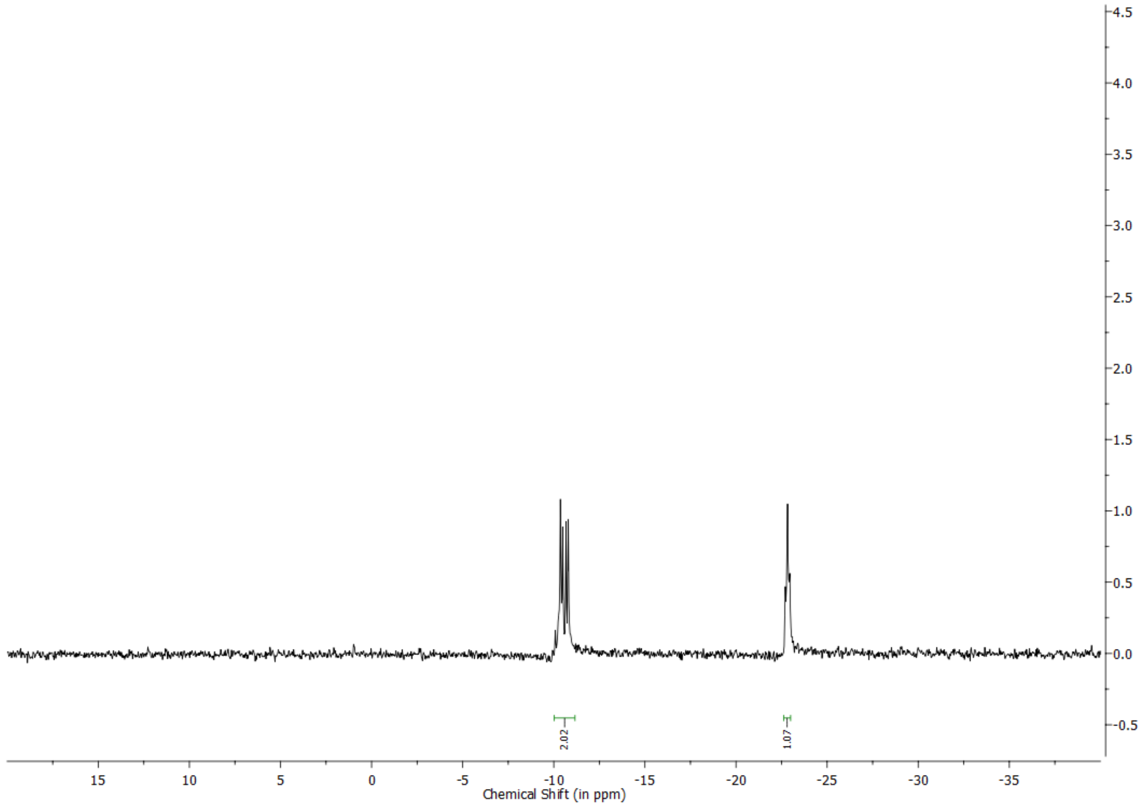


**Fig S3** 31P-NMR of prebiotic genetic alphabet BaTP in $D_2O$

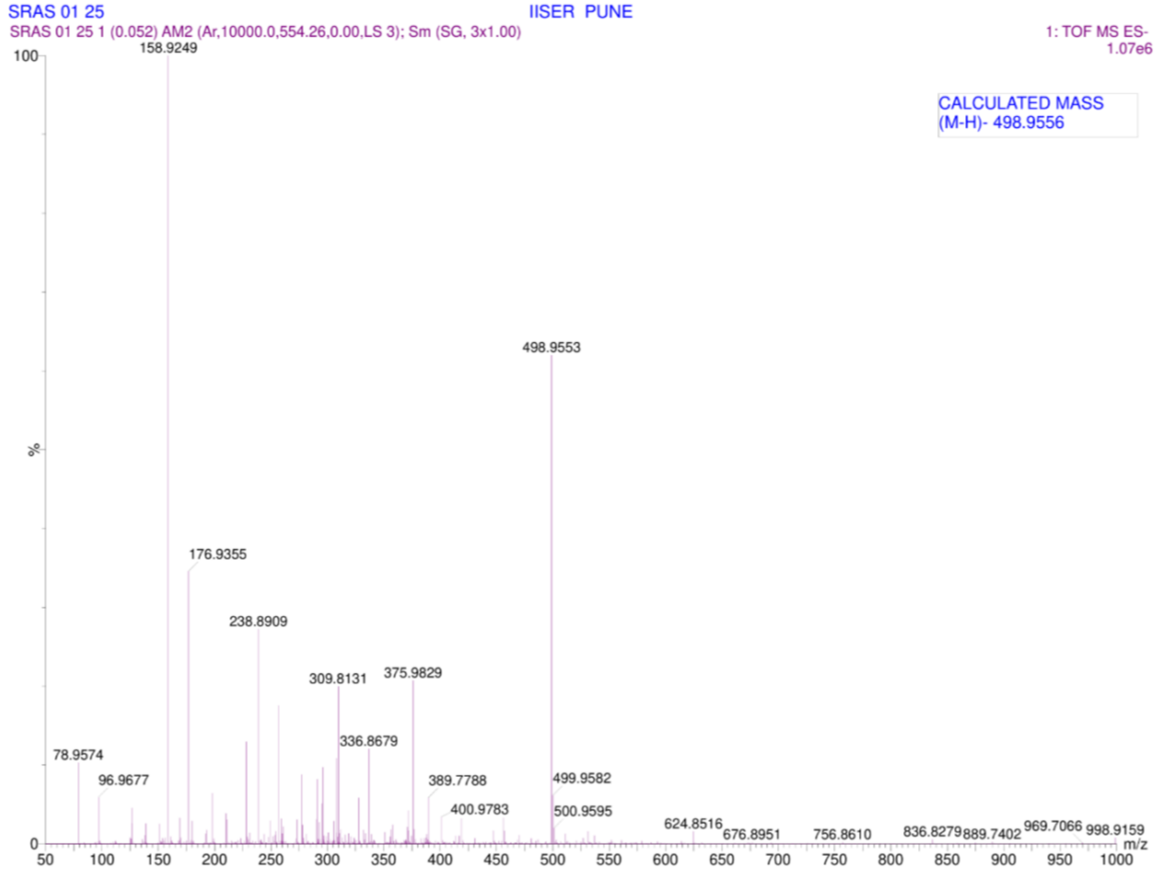


**Fig S4** HRMS of prebiotic genetic alphabet BaTP in $D_2O$

**2. Enzymatic digestion of barbitudine-modified mCherry RNA**

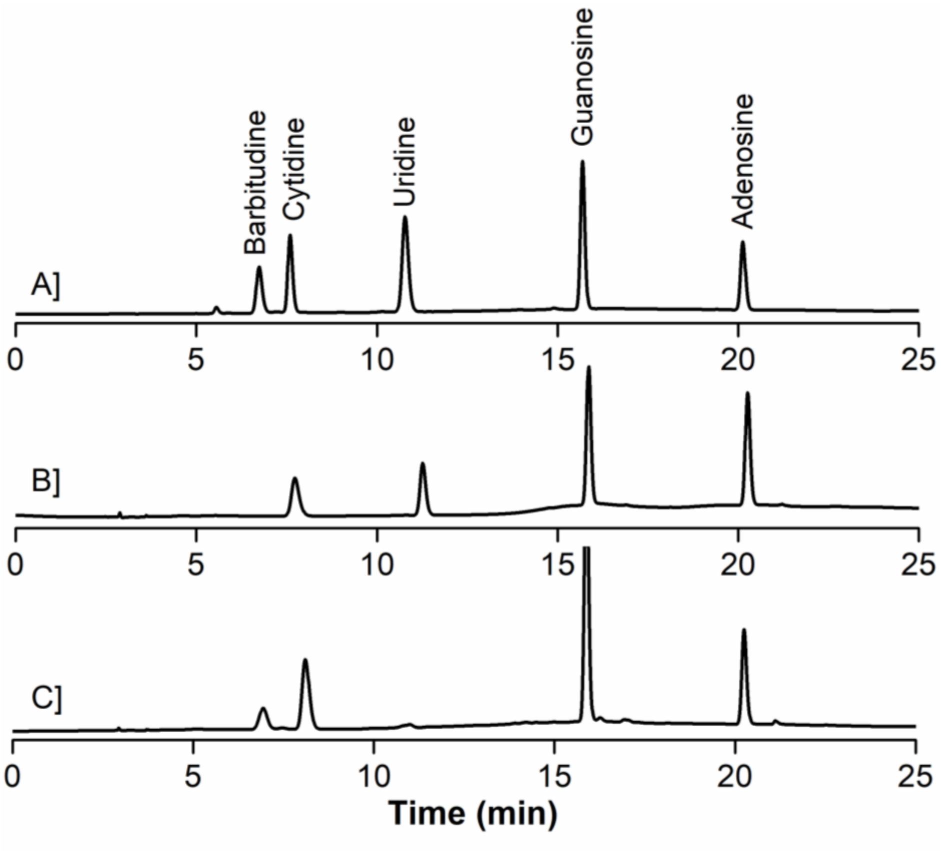


**Fig S5** HPLC chromatograms of ribonucleoside products obtained from enzymatic digestion of RNA transcripts at 260 nm. (A) Standard mix of natural ribonucleosides and Barbitudine. (B) RNA digests that were obtained from IVT by transcription reactions carried out in the presence of UTP. (C) RNA digest is obtained from IVT by transcription reaction carried out in the presence of BaTP. HPLC details - Mobile phase A: 100 mM TEAA (pH 7.5); mobile phase B: acetonitrile. Flow rate: 1 mL/min. Gradient: 0−10% B in 20 minutes and 10−100% B in 10 minutes.

#### 3. PCR product Sequencing chromatogram and sequence alignment with DNA template

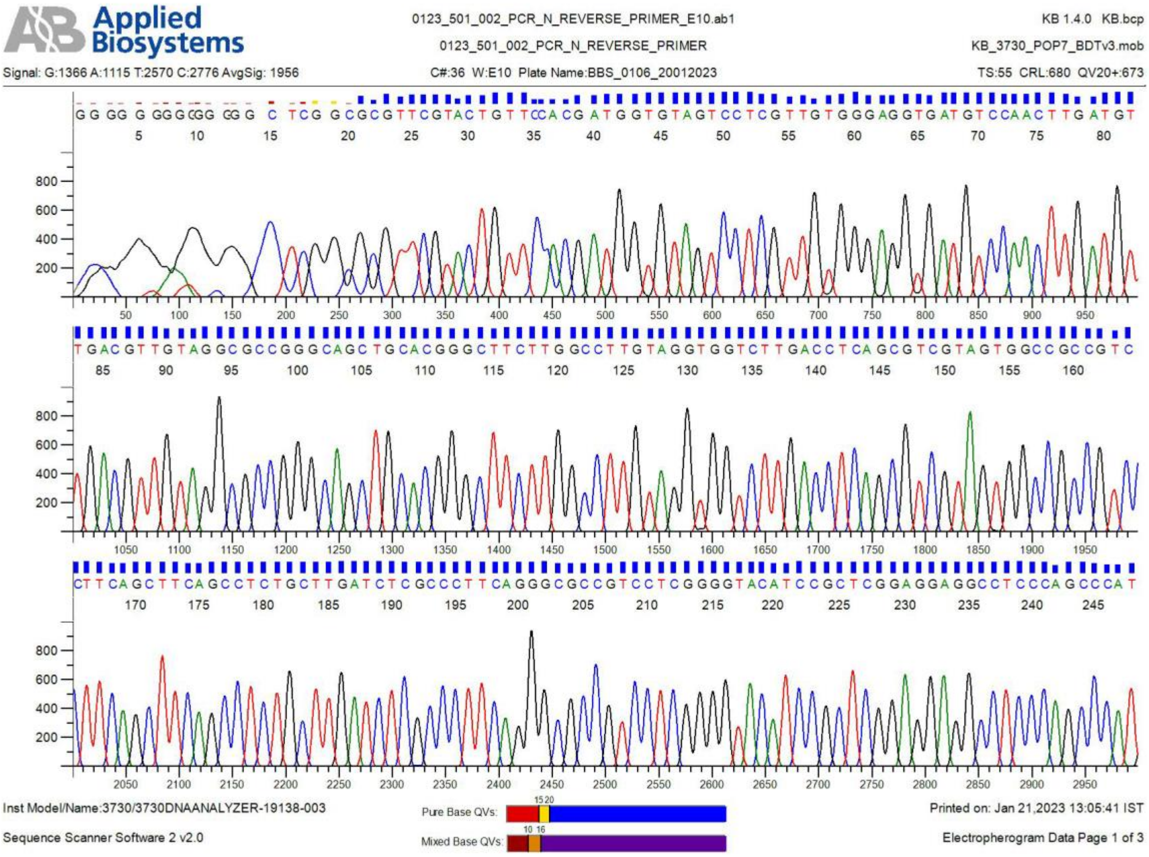


**Fig S6** Representative sequencing chromatogram of PCR amplified DNA products obtained from cDNA, which was in turn obtained by reverse transcribing control unmodified RNA. The sequencing data of the PCR product 100% sequence identities (corroborated with sequence blast Fig S8) with the template DNA sequence used for the incorporation of UTP into RNA by in vitro transcription reactions.

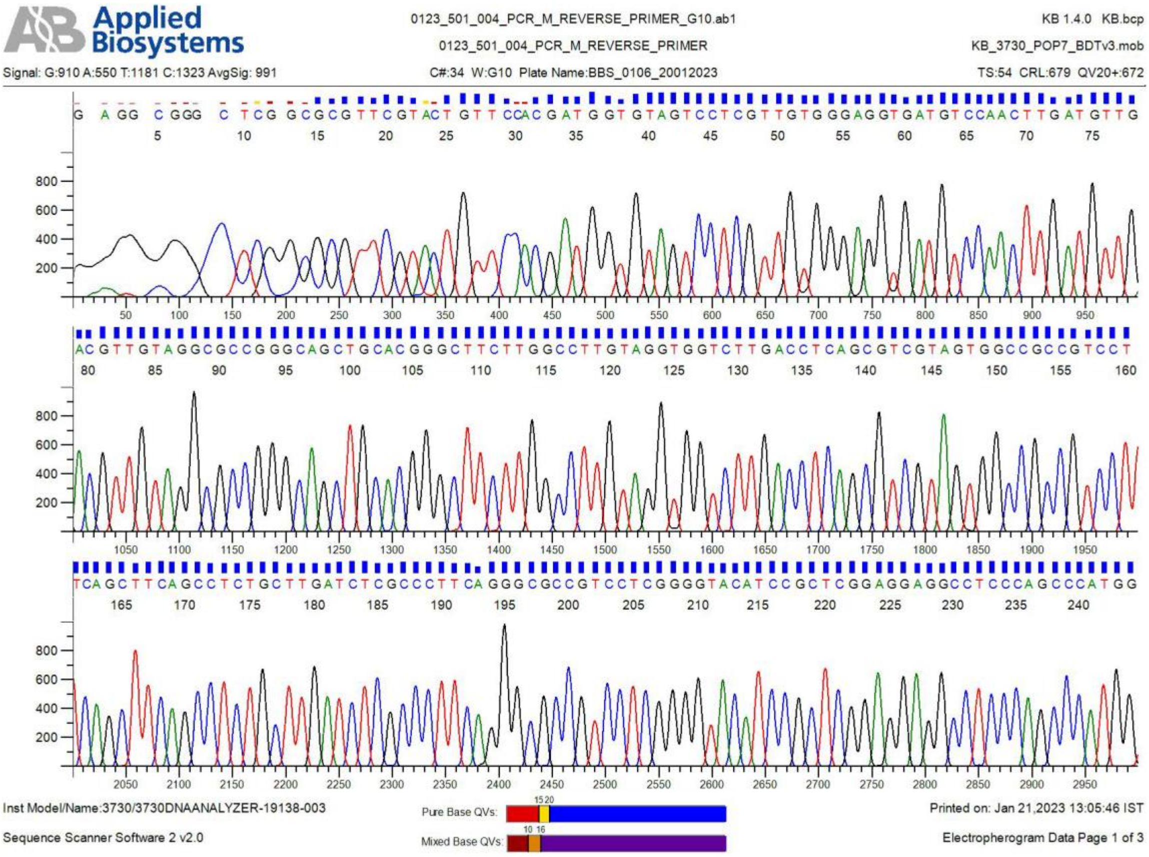


**Fig S7** Representative sequence chromatogram of PCR amplified DNA products obtained from cDNA, which was in turn obtained by reverse transcribing modified RNA containing barbitudine. The sequencing data of the PCR product showed 99 % sequence identities with the template DNA sequence used for the incorporation of BaTP into RNA by *in vitro* transcription reactions.

Download Graphics

Sequence ID: **Query_44595** Length: **708** Number of Matches: **1**

**Range 1: 15 to 683** Graphics

Next Match Pr

| Score | Expect | Identities | Gaps | Strand |
|---|---|---|---|---|
| 1236 bits(669) | 0.0 | 669/669(100%) | 0/669(0%) | Plus/Minus |

```
Query  1    ATGGTGAGCAAGGGCGAGGAGGATAACATGGCCATCATCAAGGAGTTCATGCGCTTCAAG  60
            ||||||||||||||||||||||||||||||||||||||||||||||||||||||||||||
Sbjct  683  ATGGTGAGCAAGGGCGAGGAGGATAACATGGCCATCATCAAGGAGTTCATGCGCTTCAAG  624

Query  61   GTGCACATGGAGGGCTCCGTGAACGGCCACGAGTTCGAGATCGAGGGCGAGGGCGAGGGC  120
            ||||||||||||||||||||||||||||||||||||||||||||||||||||||||||||
Sbjct  623  GTGCACATGGAGGGCTCCGTGAACGGCCACGAGTTCGAGATCGAGGGCGAGGGCGAGGGC  564

Query  121  CGCCCCTACGAGGGCACCCAGACCGCCAAGCTGAAGGTGACCAAGGGTGGCCCCCTGCCC  180
            ||||||||||||||||||||||||||||||||||||||||||||||||||||||||||||
Sbjct  563  CGCCCCTACGAGGGCACCCAGACCGCCAAGCTGAAGGTGACCAAGGGTGGCCCCCTGCCC  504

Query  181  TTCGCCTGGGACATCCTGTCCCCTCAGTTCATGTACGGCTCCAAGGCCTACGTGAAGCAC  240
            ||||||||||||||||||||||||||||||||||||||||||||||||||||||||||||
Sbjct  503  TTCGCCTGGGACATCCTGTCCCCTCAGTTCATGTACGGCTCCAAGGCCTACGTGAAGCAC  444

Query  241  CCCGCCGACATCCCCGACTACTTGAAGCTGTCCTTCCCCGAGGGCTTCAAGTGGGAGCGC  300
            ||||||||||||||||||||||||||||||||||||||||||||||||||||||||||||
Sbjct  443  CCCGCCGACATCCCCGACTACTTGAAGCTGTCCTTCCCCGAGGGCTTCAAGTGGGAGCGC  384

Query  301  GTGATGAACTTCGAGGACGGCGGCGTGGTGACCGTGACCCAGGACTCCTCCCTGCAGGAC  360
            ||||||||||||||||||||||||||||||||||||||||||||||||||||||||||||
Sbjct  383  GTGATGAACTTCGAGGACGGCGGCGTGGTGACCGTGACCCAGGACTCCTCCCTGCAGGAC  324

Query  361  GGCGAGTTCATCTACAAGGTGAAGCTGCGCGGCACCAACTTCCCCTCCGACGGCCCCGTA  420
            ||||||||||||||||||||||||||||||||||||||||||||||||||||||||||||
Sbjct  323  GGCGAGTTCATCTACAAGGTGAAGCTGCGCGGCACCAACTTCCCCTCCGACGGCCCCGTA  264

Query  421  ATGCAGAAGAAGACCATGGGCTGGGAGGCCTCCTCCGAGCGGATGTACCCCGAGGACGGC  480
            ||||||||||||||||||||||||||||||||||||||||||||||||||||||||||||
Sbjct  263  ATGCAGAAGAAGACCATGGGCTGGGAGGCCTCCTCCGAGCGGATGTACCCCGAGGACGGC  204

Query  481  GCCCTGAAGGGCGAGATCAAGCAGAGGCTGAAGCTGAAGGACGGCGGCCACTACGACGCT  540
            ||||||||||||||||||||||||||||||||||||||||||||||||||||||||||||
Sbjct  203  GCCCTGAAGGGCGAGATCAAGCAGAGGCTGAAGCTGAAGGACGGCGGCCACTACGACGCT  144

Query  541  GAGGTCAAGACCACCTACAAGGCCAAGAAGCCCGTGCAGCTGCCCGGCGCCTACAACGTC  600
            ||||||||||||||||||||||||||||||||||||||||||||||||||||||||||||
Sbjct  143  GAGGTCAAGACCACCTACAAGGCCAAGAAGCCCGTGCAGCTGCCCGGCGCCTACAACGTC  84

Query  601  AACATCAAGTTGGACATCACCTCCCACAACGAGGACTACACCATCGTGGAACAGTACGAA  660
            ||||||||||||||||||||||||||||||||||||||||||||||||||||||||||||
Sbjct  83   AACATCAAGTTGGACATCACCTCCCACAACGAGGACTACACCATCGTGGAACAGTACGAA  24

Query  661  CGCGCCGAG  669
            |||||||||
Sbjct  23   CGCGCCGAG  15
```

**Fig S8** Representative sequence alignment (BLAST) of PCR amplified DNA product obtained from cDNA, which was in turn obtained by reverse transcribing unmodified RNA. The sequencing data of the PCR product showed 100% sequence identities (corroborated with sequence blast Fig S9) with the template DNA sequence used for the incorporation of UTP into RNA by in vitro transcription reactions. Importantly, sequence blast clearly showed that no misincorporation was observed while incorporating thymidine against adenosine residue in the cDNA template.

Download Graphics

Sequence ID: **Query_27161** Length: **702** Number of Matches: **1**

**Range 1: 3 to 677** Graphics    ▼ Next Match ▲ P

| Score | Expect | Identities | Gaps | Strand |
|---|---|---|---|---|
| 1236 bits(669) | 0.0 | 674/676(99%) | 1/676(0%) | Plus/Minus |

```
Query  1    ATGGTGAGCAAGGGCGAGGAGGATAACATGGCCATCATCAAGGAGTTCATGCGCTTCAAG  60
            ||||||||||||||||||||||||||||||||||||||||||||||||||||||||||||
Sbjct  677  ATGGTGAGCAAGGGCGAGGAGGATAACATGGCCATCATCAAGGAGTTCATGCGCTTCAAG  618

Query  61   GTGCACATGGAGGGCTCCGTGAACGGCCACGAGTTCGAGATCGAGGGCGAGGGCGAGGGC  120
            ||||||||||||||||||||||||||||||||||||||||||||||||||||||||||||
Sbjct  617  GTGCACATGGAGGGCTCCGTGAACGGCCACGAGTTCGAGATCGAGGGCGAGGGCGAGGGC  558

Query  121  CGCCCCTACGAGGGCACCCAGACCGCCAAGCTGAAGGTGACCAAGGGTGGCCCCCTGCCC  180
            ||||||||||||||||||||||||||||||||||||||||||||||||||||||||||||
Sbjct  557  CGCCCCTACGAGGGCACCCAGACCGCCAAGCTGAAGGTGACCAAGGGTGGCCCCCTGCCC  498

Query  181  TTCGCCTGGGACATCCTGTCCCCTCAGTTCATGTACGGCTCCAAGGCCTACGTGAAGCAC  240
            ||||||||||||||||||||||||||||||||||||||||||||||||||||||||||||
Sbjct  497  TTCGCCTGGGACATCCTGTCCCCTCAGTTCATGTACGGCTCCAAGGCCTACGTGAAGCAC  438

Query  241  CCCGCCGACATCCCCGACTACTTGAAGCTGTCCTTCCCCGAGGGCTTCAAGTGGGAGCGC  300
            ||||||||||||||||||||||||||||||||||||||||||||||||||||||||||||
Sbjct  437  CCCGCCGACATCCCCGACTACTTGAAGCTGTCCTTCCCCGAGGGCTTCAAGTGGGAGCGC  378

Query  301  GTGATGAACTTCGAGGACGGCGGCGTGGTGACCGTGACCCAGGACTCCTCCCTGCAGGAC  360
            ||||||||||||||||||||||||||||||||||||||||||||||||||||||||||||
Sbjct  377  GTGATGAACTTCGAGGACGGCGGCGTGGTGACCGTGACCCAGGACTCCTCCCTGCAGGAC  318

Query  361  GGCGAGTTCATCTACAAGGTGAAGCTGCGCGGCACCAACTTCCCCTCCGACGGCCCCGTA  420
            ||||||||||||||||||||||||||||||||||||||||||||||||||||||||||||
Sbjct  317  GGCGAGTTCATCTACAAGGTGAAGCTGCGCGGCACCAACTTCCCCTCCGACGGCCCCGTA  258

Query  421  ATGCAGAAGAAGACCATGGGCTGGGAGGCCTCCTCCGAGCGGATGTACCCCGAGGACGGC  480
            ||||||||||||||||||||||||||||||||||||||||||||||||||||||||||||
Sbjct  257  ATGCAGAAGAAGACCATGGGCTGGGAGGCCTCCTCCGAGCGGATGTACCCCGAGGACGGC  198

Query  481  GCCCTGAAGGGCGAGATCAAGCAGAGGCTGAAGCTGAAGGACGGCGGCCACTACGACGCT  540
            ||||||||||||||||||||||||||||||||||||||||||||||||||||||||||||
Sbjct  197  GCCCTGAAGGGCGAGATCAAGCAGAGGCTGAAGCTGAAGGACGGCGGCCACTACGACGCT  138

Query  541  GAGGTCAAGACCACCTACAAGGCCAAGAAGCCCGTGCAGCTGCCCGGCGCCTACAACGTC  600
            ||||||||||||||||||||||||||||||||||||||||||||||||||||||||||||
Sbjct  137  GAGGTCAAGACCACCTACAAGGCCAAGAAGCCCGTGCAGCTGCCCGGCGCCTACAACGTC  78

Query  601  AACATCAAGTTGGACATCACCTCCCACAACGAGGACTACACCATCGTGGAACAGTACGAA  660
            ||||||||||||||||||||||||||||||||||||||||||||||||||||||||||||
Sbjct  77   AACATCAAGTTGGACATCACCTCCCACAACGAGGACTACACCATCGTGGAACAGTACGAA  18

Query  661  CGCGCCGAGGGCCGCC  676
            |||||||||  |||||
Sbjct  17   CGCGCCGAGC-CCGCC  3
```

**Fig S9** Representative sequence alignment (BLAST) of PCR amplified DNA products obtained from cDNA, which was in turn obtained by reverse transcribing modified RNA containing barbitudine. The sequencing data of the PCR product 99 % sequence identities with the template DNA sequence used for the incorporation of BaTP into RNA by *in vitro* transcription reactions. Importantly, sequence blast clearly showed that no misincorporation was observed while incorporating thymidine against adenosine residue in the cDNA template.
